# KSHV Terminal Repeat Regulates Inducible Lytic Gene Promoters

**DOI:** 10.1101/2023.09.07.556745

**Authors:** Yoshihiro Izumiya, Adhraa Algalil, Jonna M. Espera, Hiroki Miura, Tomoki Inagaki, Chie Izumiya, Ashish Kumar

## Abstract

The Kaposi’s sarcoma-associated herpesvirus (KSHV) genome consists of an approximately 140 kb unique coding region flanked by multiple copies of 0.8 kb terminal repeat (TR) sequence. While TR’s function in plasmid maintenance is well-established, TR’s transcription regulatory roles have not been fully explored. Here, we show KSHV TR is a large transcription regulatory domain.

A series of Cleavage Under Targets & Release Using Nuclease demonstrated that TR fragments are occupied by histone modifying enzymes that are known to interact with LANA in naturally infected cells, and the TR possessed characteristic enhancer histone modifications. The H3K4me3 and H3K27Ac modifications were conserved in unique region of the KSHV genome among naturally infected cells, and the KSHV Origin of lytic replication (Ori-Lyt) showed similar protein and histone modification occupancies with TR’s. In the Ori-Lyt region, the LANA complex colocalizes with H3K27Ac-modified nucleosome along with paused RNA polymerase II, and two K-Rta recruitment sites frank the nucleosome. The isolated reporter assays demonstrated that neighboring TR fragments enhanced viral lytic gene promoter activity independent of orientation in KSHV-infected and non-infected 293FT cells. K-Rta transactivation function was drastically enhanced with TR, while LANA acquired promoter repression function with TR. KSHV TR is, therefore a regulatory domain for KSHV inducible genes. However, in contrast to cellular enhancers that are bound by multiple transcription factors, perhaps the KSHV enhancer is predominantly regulated by the LANA nuclear body with TR. KSHV evolved a clever mechanism to tightly control the latency-lytic switch with the TR/LANA complex.

**Importance:** Enhancers are a crucial regulator of differential gene expression programs. Enhancer is the cis-regulatory sequences that determine target genes’ spatiotemporal and quantitative expression. Here, we show that KSHV terminal repeats fulfill the enhancer definition for KSHV inducible gene promoters. KSHV enhancer is occupied by LANA and its interacting proteins, such as CHD4, and CHD4 is known to restrict enhancers to access promoters for activation. This study thus proposes a new latency-lytic switch model in which TR accessibility to the KSHV gene promoters regulates lytic gene transcription.

## Introduction

There is growing awareness of the need to understand the spatial and temporal organization of the genome and its role in regulating gene expression. Given that nuclear enzymes and transcriptional factors are in limited supply and cannot be everywhere in the nucleus at the same time, the nuclear architecture (including chromatin structure and protein distribution) must allow for these molecules to be concentrated at the proper time and place for coordinated gene expression to occur. Understanding the three-dimensional (3D) structure of the genome and its impact on neighboring genomic elements is therefore critical to understanding gene regulation(1–5). The significance is also highlighted by the fact that deregulation of genomic interactions by mutations of intragenic genomic regions and nuclear remodeling factors such as SWI/SNF (SWItch/Sucrose Non-Fermentable) frequently leads to diseases (6, 7).

With assisted by the three-dimensional genomic architecture, gene enhancers are known to act in *cis* and in an orientation-independent fashion by forming genomic loops to increase frequencies for transcription initiation at promoters. By forming genomic looping with multiple promoters, the enhancer DNA fragment neighbors to and activates many promoters at the same time(8, 9). A larger genomic region collectively bound by an array of transcription factors at higher density, hence harboring a higher density of transcription enzymes, is called a super-enhancer(10). Cellular proteins, such as mediator complex subunit 1 (MED1) and bromodomain-containing protein 4 (BRD4), are known to be located at the super enhancer region, and those proteins further compartmentalize nuclear condensates with their intrinsically disordered domain for maintaining selective gene expression hence cell-identity(11). The genomic interaction between enhancer and promoter mediated by transcription-related enzymes positions RNA Pol II pre-initiation complex at specific genomic sites for sensitizing inducible gene expression with specific signaling events that activate enhancers(2, 11–16).

The KSHV viral genome consists of an approximately 140 kb unique coding region flanked by multiple TRs of 801 bp with high GC content(17). KSHV genomes persist in latently infected cells as episomes via tethering to the host cell chromosomes. During latency, the expression of lytic viral genes is poised to be transcribed, and only a few latent genes are actively expressed and translated(18). Among these latent genes, ORF73 encodes latency-associated nuclear antigen (LANA), which plays a crucial role in latent episomal DNA replication and segregation during host cell mitosis. The TR contains a DNA replication origin called ori-P that consists of two LANA-biding sites (LBS): a higher affinity site (LBS1) and a lower affinity site (LBS2) followed by an adjacent 32-bp GC-rich segment(18). Episome maintenance requires at least two LBS1/2 binding sites, and the viral genome consists of 30-40 TRs(17, 19). A crystal structure demonstrated that LANA_DBD_ mainly exists as a dimer in solution, and five LANA_DBD_ dimers can interact end-to-end to form a decameric ring with an exterior diameter of 110 Å and an interior diameter of 50 Å, and the inner diameter accommodates double-stranded DNA(20). DNA binding further induces oligomerization of LANA_DBD_, and a hydrophobic interface between LANA dimer to form the decameric ring is essential for cooperative DNA binding and DNA replication, hence episome maintenance (20). The specific LANA’s TR DNA binding also increases up to 600-800 (2×5×2×[30-40]) LANA protein copies to locate on a single KSHV episome. As the result, LANA can be seen as LANA dots in KSHV-infected cells with immunostaining (21). Assist from KSHV 3D genomic structure with episomal genomes (22–25), the TRs always localize relatively close proximity to unique region that encode many lytic gene promoters, and those promoters are known to be activated by <u>K</u>SHV <u>R</u>eplication and <u>T</u>ranscription <u>A</u>ctivator (K-Rta) protein. Isolated reporter constructs showed that at least 33 lytic gene promoters can be activated by K-Rta in 293 cells (26). While TR’s function in plasmid maintenance is well-established, TR function in transcription regulation remains unclear, except a publication that demonstrated that K1 gene promoter activity is enhanced by including partial TR sequence (27).

Here, we show that TR fulfills the definition of gene enhancer for KSHV lytic gene promoters and is a transcription regulatory element for KSHV lytic gene promoters. With reporter assays, we demonstrated that TR possesses orientation-independent transactivation function. We also show that previously identified LANA interacting transcription factors are accumulated at the TR region in naturally infected cells. Having TR in the reporter constructs substantially enhanced K-Rta and LANA transcription function. Here, we propose a KSHV gene regulatory model in which KSHV TR is a large transcription regulatory domain for KSHV promoters (unique region).

## Results

### Latent KSHV Chromatin Modification Maps

To understand the KSHV latency-lytic switch, we first revisited a comprehensive histone modification of the KSHV episomes in naturally infected cells. We used a public database and our CUT&RUN (Cleavage Under Targets & Release Using Nuclease) data sets. If having a specific regulatory genomic domain is necessary for maintaining inducible KSHV latent chromatins, cell lines that can produce infections virions with stimulation should conserve similar genomic domains in KSHV latent chromatin. Four histone modifications primarily associated with active transcription (H3K4me1, H3K4me2, H3K4me3, and H3K27Ac), and repressive marks (H3K27me3, H3K9me3) were examined. While we tried to locate H3K9me3, we could not identify reasonable peaks with two independent antibodies.

Figure 1A depicts histone modifications of KSHV latent chromatin in BCBL-1 cells. The results showed that four active histone marks are clustered mainly at genomic loci encoding early and immediate-early genes consistent with previous reports(28, 29). The H3K27Ac marks are restricted at two Ori-Lyt regions, PAN promoter, vIRF coding loci, LANA promoter, ORF75 promoter, and K15 promoter regions (Fig. 1A). Although H3K4me1 and me2 marks are co-localized with H3K4me3 marks, farmer two modifications were more broadly distributed around the H3K4me3 marks. The H3K4me3 modification showed sharper and stronger signals (higher peaks) than H3K4me1 and me2 marks in BCBL-1 (Fig. 1A). While it is difficult to compare between peaks at the unique genomic region of the KSHV genome (one copy per genome) and that of terminal repeat fragments (20-40 copies per genome), there are still noticeably strong H3K4me3 and H3K27Ac signals at the TR regions but not with the H3K4me1, 2, or H3K27me3 (Fig. 1A **right panel**). On the other hand, repressive marks (H3K27me3) were localized more broadly in BCBL-1. Other KSHV naturally infected PEL cells or experimentally-infected iSLK cells showed similar histone modification patterns for active histone marks, especially for H3K4me3 and H3K27Ac marks. However, H3K27me3 had different degrees of signals at late gene cluster regions. With that, BC3 had strongest signals, while BC-1 cells showed little signals at the late gene cluster regions (Fig. 1B). The experimentally-infected iSLK cells also showed patterns more to resembling BC3 and globally occupied by H3K27me3 modified histones. These results indicate that maintaining active genomic regions is a more conserved trait than the repressive mark for KSHV latent chromatin.

**Figure 1.**
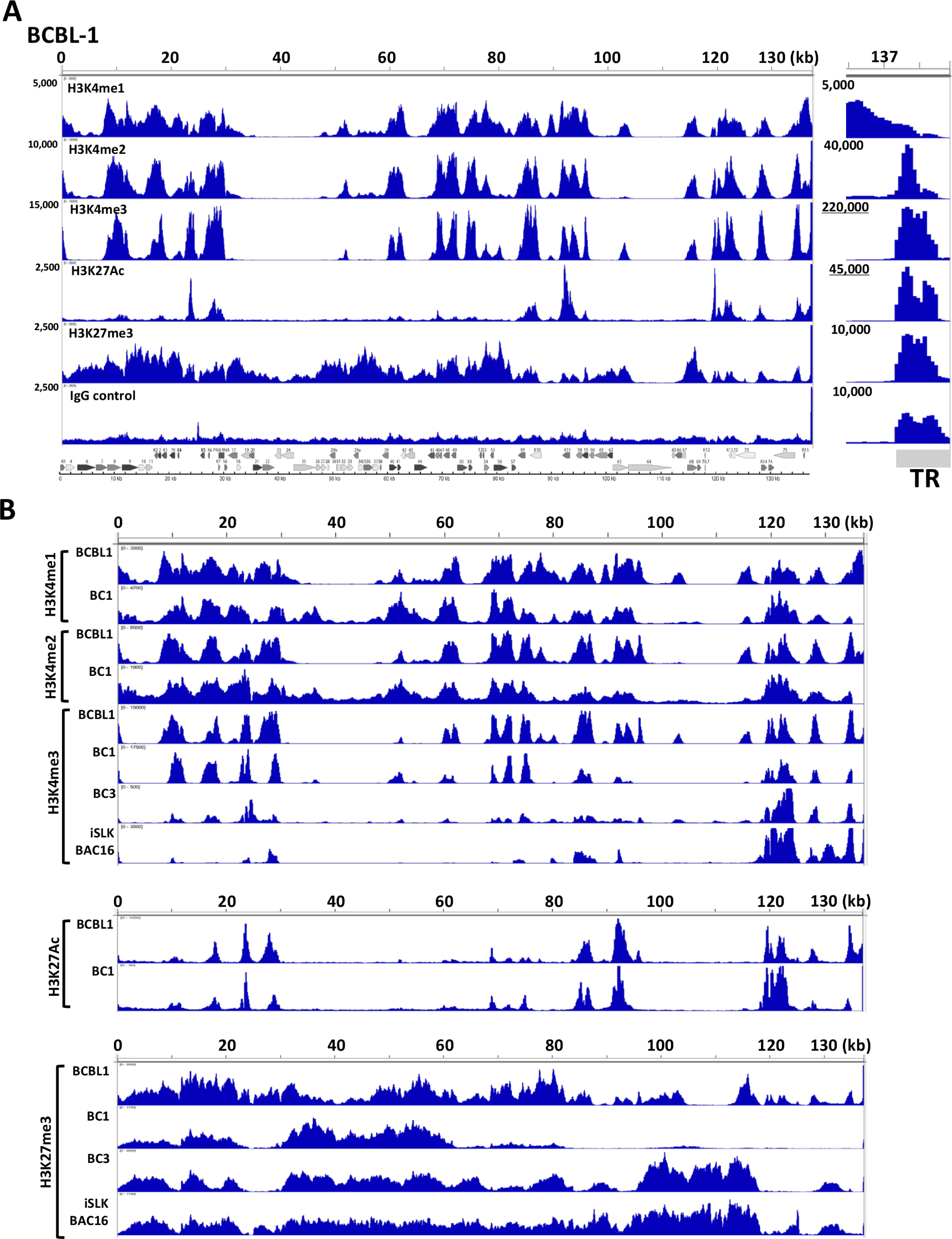
KSHV chromatin modifications. (A) KSHV latent chromatin modifications in BCBL-1. The position of indicated histone modification was examined with CUT&RUN analyses. The respective peak heights are indicated on the left panel. KSHV ORF map was depicted at the bottom of the panel. The terminal repeat region is zoomed at right after adjusting the scale of peak ranges. Sequence enrichments seen more than 10 times compared with that of the unique region was underlined. **(B) Differential chromatin modifications in KSHV infected cells.** Selected histone modifications were compared among KSHV infected cells. Cell line name was depicted in the left of the panel. The H3K27Ac ChIP-seq data for BC1 and BC3 are adapted from SRR9956027 and SRR9956035, respectively.

### Protein interaction among K-Rta and LANA interacting proteins

To have insights into the latency-lytic switch mechanism, we next focused on KSHV LANA and K-Rta interacting transcription-related enzymes. This is because enzymes regulate the chromatin modification and occupancies. KSHV LANA is essential for establishing and maintaining latent infection, and we previously identified proteins localized in proximity to LANA (24). Among the LANA interacting proteins (Fig. 2 **blue** oval), we showed that cellular CHD4 protein plays a key role in establishing and maintaining latent infection (24). The proximity-biotin labeling with recombinant KSHV also confirmed that LANA interacts with BRD4, and also identified components of I-SWI/SNF complex, and several others as LANA interacting proteins (Fig. 2 blue oval (24)). In addition, to reveal how K-Rta triggers KSHV reactivation, we also reported cellular proteins that are induced to interact with RNAPII and K-Rta protein during KSHV reactivation. In the latter study, we applied rapid immunoprecipitation mass spectrometry of endogenous protein (RIME) (30), and the method is suitable for identifying transcription factor complexes on chromatin. By identifying proteins that interact RNAPII and K-Rta during reactivation on chromatin, we isolated critical proteins for K-Rta transactivation (30). The study identified that K-Rta interacted with NCoA2, SWI/SNF, and Mediator complex, and these proteins may form a large protein complex (Fig. 2 red oval). The results were consistent with previous reports that showed K-Rta interaction with NCoA2 and SWI/SNF and Mediators (31, 32). To provide insights into dynamic protein interactions between two protein complexes during reactivation, we combined and visualized putative protein interactions among the two protein complexes. The result showed that LANA interacting proteins (blue ovals) can interact with components of the K-Rta protein complex (red ovals), when they were recruited to proximity during reactivation. Nearly all, except two (SMCHD1 and HP1BP3) of components of LANA interacting proteins, can bind to at least one of the components of the K-Rta protein complex (Fig. 2). The protein interaction model suggests that components of the LANA protein complex at TR (during latency) can be flexible when K-Rta brought its protein complex near the LANA binding sites during reactivation.

**Figure 2.**
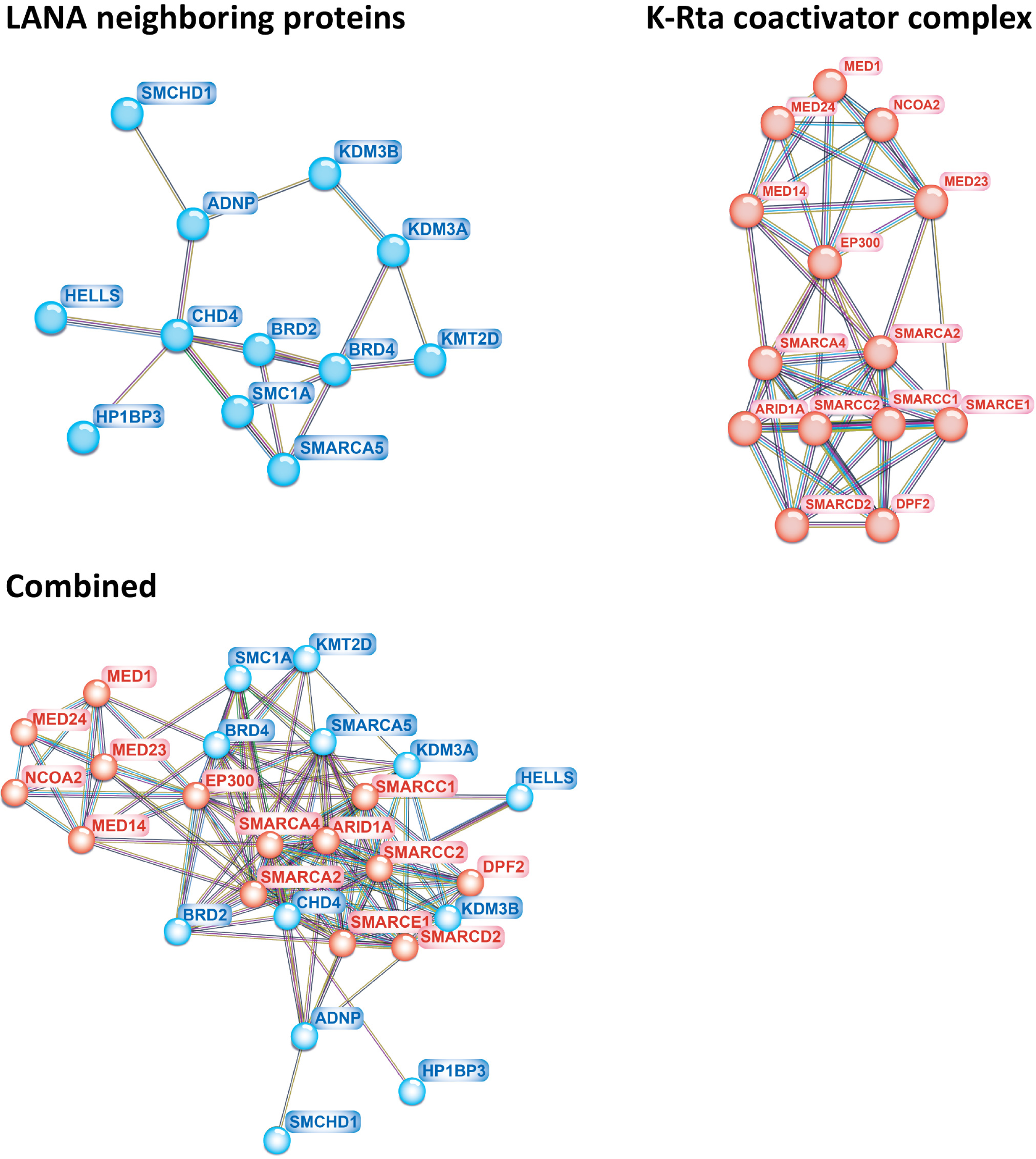
LANA and K-Rta protein complexes. (A) LANA neighboring protein in iSLK cells. Previously identified LANA neighboring chromatin binding proteins by proximity biotin labeling with recombinant KSHV were visualized with STRING. **(B) K-Rta and RNA polymerase II interacting protein on chromatin.** Previously identified K-Rta interacting proteins by RIME study were visualized with STRING. The RIME showed formation of the putative protein complex. **(C) Putative protein interactions among LANA and K-Rta complexes.** Theoretical protein complex interactions in the presence of K-Rta are simulated with STRING.

### Determining LANA interacting protein recruitment sites on KSHV genome

To gain insights for the gene regulation, we next examined occupancies of the LANA interacting enzymes by CUT&RUN in latently infected cells. We selected targeting molecules based on the availability of antibodies that have been proven for successful CUT&RUNs. The results showed that LANA interacting enzymes and cofactors (e.g., BRD4, KMT2D, SMARCA5, CHD4) are colocalized with KSHV LANA (Fig. 3A). As expected, the LANA interacting proteins were enriched at TR region. The scale of the integrated genome browser was adjusted with x20 of the unique region to consider multiple TR copies. The LANA sequence reads at TR were still off the chart, and the highest LANA peaks demonstrated over 6,000,000 sequence reads. No comparable signals were seen with any other proteins tested or genomic regions. Because peaks at TR for the CTCF, H3K27me3, or H3K4me1 were lower than sequence reads found in the unique region, we considered that generally higher CUT&RUN signals at TR regions are unlikely due to backgrounds. Accordingly, we considered that the presence of comparable peaks with unique region after scaling changed to the x20 (Fig. 3A, right panel) as a presence at TR. With the criteria, we suggest that, at least, RNAPII, LANA, BRD4, SMARCA5, CHD4, and ADNP localizes at TR during latency in BCBL-1. The lack of strong and sharp peaks for CTCF or SMC1 also suggested no cohesin-mediated "fixed" genomic loop formations with TR.

**Figure 3.**
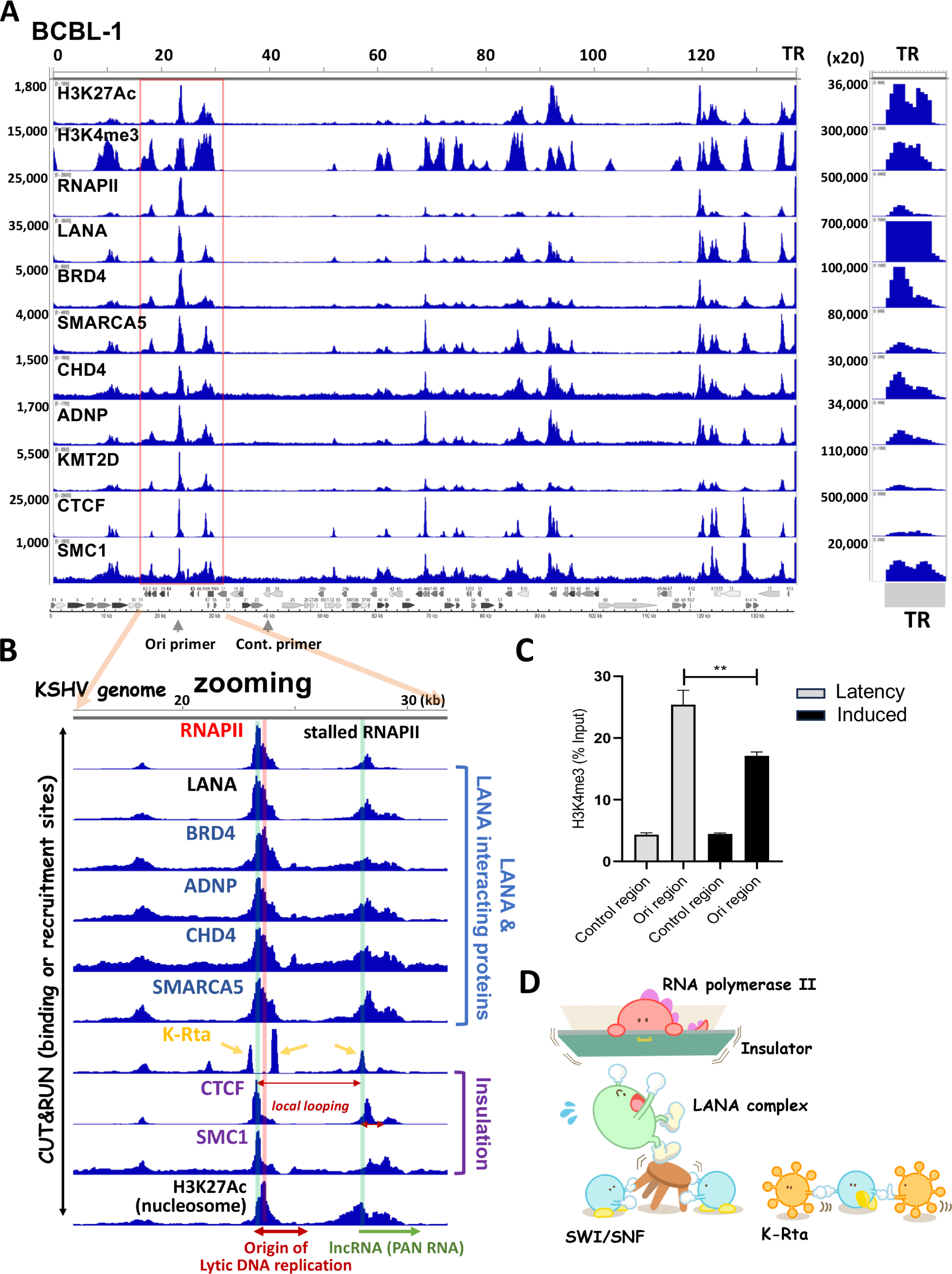
Histone enzyme recruitment sites. (A) Overview of LANA complex recruitment sites. LANA interacting protein recruitment sites were determined by CUT&RUN and mapped on the KSHV genome. The range of sequence reads is shown on the left of the panel, TR region was zoomed at right and peak scale was adjusted by multiplying by 20 to consider the TR copy number. The KSHV ORF map was depicted at the bottom of the panel. Selective LANA interacting transcription-related proteins are accumulated at TR. Position of primers used for (C) was indicated at the bottom of the panel. **(B) Zooming at Ori-Lyt and PAN RNA promoter in the unique region.** One of the LANA complexes recruited sites at the unique region was zoomed, and positional association among CTCF/SMC1 (insulation), H3K27Ac nucleosome, LANA complex recruitment site, and K-Rta binding sites are emphasized. The position of the nucleosome (red shadow) and CTCF binding sites (green shadow) are also indicated. Association with the frequencies of genomic looing is depicted in supplemental figure 1. The origin of Lyic DNA replication and PAN RNA regions are highlighted at the bottom of the panel. **(C) The nucleosome eviction at the Origin of DNA replication.** TREx-K-Rta BCBL-1 cells were left untreated or treated with Doxycycline (1 μg/ml) and TPA (20 ng/ml) for 48 h. Cut & Run was performed using H3K4me3 antibody. The percent of input was calculated and shown in the panel. H3K4me3 non-binding region (Control region) was used as a negative control. Unpaired t-test was used for calculating the p-value. **(D) Proposing Latency-Lytic switch model.** Through sequence-specific DNA binding, K-Rta recruits SWI/SNF complex at K-Rta binding sites. The SWI/SNF complex slides/evicts nucleosomes that the LANA complex is tethering. LANA complex might bind the H3K27Ac through direct histone binding or BRD4. The destabilization of LANA complex and nucleosome eviction facilitates RNAPII elongation, which is stalled at Ori-Lyt and PAN RNA promoter regions.

Even though transcription factors presented in Fig. 3 were found to be localized in close proximity to LANA, occupancies of those enzymes at TR (height of peaks at unique region/ height of peaks at TR for each LANA interacting protein) were not identical. The results may suggest either highly heterogenic recruitment at an episomal level in an infected cell or dynamic protein recruitment at the individual TR copy. If the all TR fragment is equally bound by the protein via LANA, we would expect peaks similar to LANA.

### Nucleosome organization at Origin of Lytic DNA replication

The observation, which LANA interacting proteins (transcription factor complex for latent maintenance) occupied at epigenetically active unique regions and the fact that viral latent chromatins are not actively transcribed, suggested that regulation of transcription initiation at the genomic region may play an important role for reactivation. Zooming the KSHV genomic region (22,000 to 29,000), encompassing Ori-Lyt and PAN RNA coding regions, where the KSHV latent chromatin possessed conserved active histone modifications (Fig. 1B), showed that LANA, CHD4, ADNP, SMARCA5, and BRD4 are recruited to next to CTCF/SMC1 peak (Fig. 3B), and the CTCF/SMC1 binding site creates a small boundary of transcription regulatory domain (23) (Fig. 3B **CTCF binding sites marked blue shadow**). The LANA tethered-H3K27Ac nucleosome localizes right next to the RNAPII peak (Fig. 3B **red shadow**), suggesting that the H3K27Ac-nucleosome may play critical role in stalling the RNAPII. Two K-Rta binding sites clearly located next to the LANA bound nucleosome. Directionality and position of CTCF/SMC1 as well as previous Hi-C sequence reads (23) suggested that Ori-Lyt forms small genomic loops with PAN RNA promoter region and the genomic loop harbors the paused RNAPII (a combined figure with previous Hi-C studies is presented in **S-**Fig. 1). The RNAPII peaks at Ori-Lyt are the strongest, when we considered TR copy number in all three PEL cell lines (Fig. 3B), suggesting that majority of infected episomes in a cell in PEL cell lines (BC-1 and BCBL-1) harbor the paused RNAPII at the same position. Furthermore, K-Rta recruitment may be designed to initiate the RNAPII elongation at the transcription regulatory domain. Because K-Rta binding sites are situated next to the LANA-bound H3K27Ac-modified nucleosome, and the K-Rta protein complex contains SWI/SNF (Fig. 2), which can evict histones to facilitate transcription elongation. Indeed, the nucleosomes occupancies were significantly decreased when we trigger reactivation with K-Rta expression (Fig. 3C). The dissociating LANA complex by eviction of LANA-BRD4 bound H3K27Ac would also regulate CTCF1/SMC1A binding, because CHD4/SMARCA5 is known to regulate CTCF1/SMC1A positioning(33–35). Based on the histone modification and protein recruitment maps, we formulated a KSHV latency-lytic switch model, in which eviction of LANA-tethering nucleosome by the K-Rta complex plays a critical role in triggering reactivation (Fig. 3D).

### Genomic interaction between TR and unique region

Having similar protein complex recruited at selected loci in unique regions and TR, we asked if these two genomic regions are located in particularly closer proximity in 3D genomic structure. To study this, we reanalyzed previous Hi-C sequence data sets with TREx-K-Rta BCBL-1 cells(23). We first isolated KSHV TR fragments and mapped the position of ligated KSHV DNA fragments onto unique region of the KSHV genome. The results showed that a significant majority of TR fragments (>99.5%) were ligated with other TR fragments (Fig. 4A), suggesting that individual TR unit is highly compacted with each other. The result agreed with studies on LANA nuclear bodies with super-resolution fluorescence microscope analyses(36). KSHV LANA (37) and BRD4 (11) are also known to form phase separate protein condensates that may further facilitates TR fragments to be compartmentalized. Even though overall frequencies are much less (< 0.5% of total TR genomic loops), two genomic regions were more frequently formed genomic loops with TR within the unique region (Fig. 4). These regions were within the Ori-PAN RNA transcription regulatory domain (**S-**Fig. 1) and near the K-Rta promoter region. Induction of KSHV reactivation by K-Rta expression increased overall KSHV-KSHV genomic loops(38), which also increased genomic interaction from TRs (Fig. 4B). The results indicated that KSHV TR DNA fragments formed self-aggregate with LANA. Considering CTCF frequently locates at epigenetically active genomic regions, having no CTCF binding sites within 25 kb+ of highly active DNA fragment (Fig. 1A) could be considered a unique genomic feature. This design would make the TR a "mobile enhancer" for KSHV promoters, because the TR fragments are maintained to be near viral promoters due to the episomal genomic structure.

**Figure 4.**
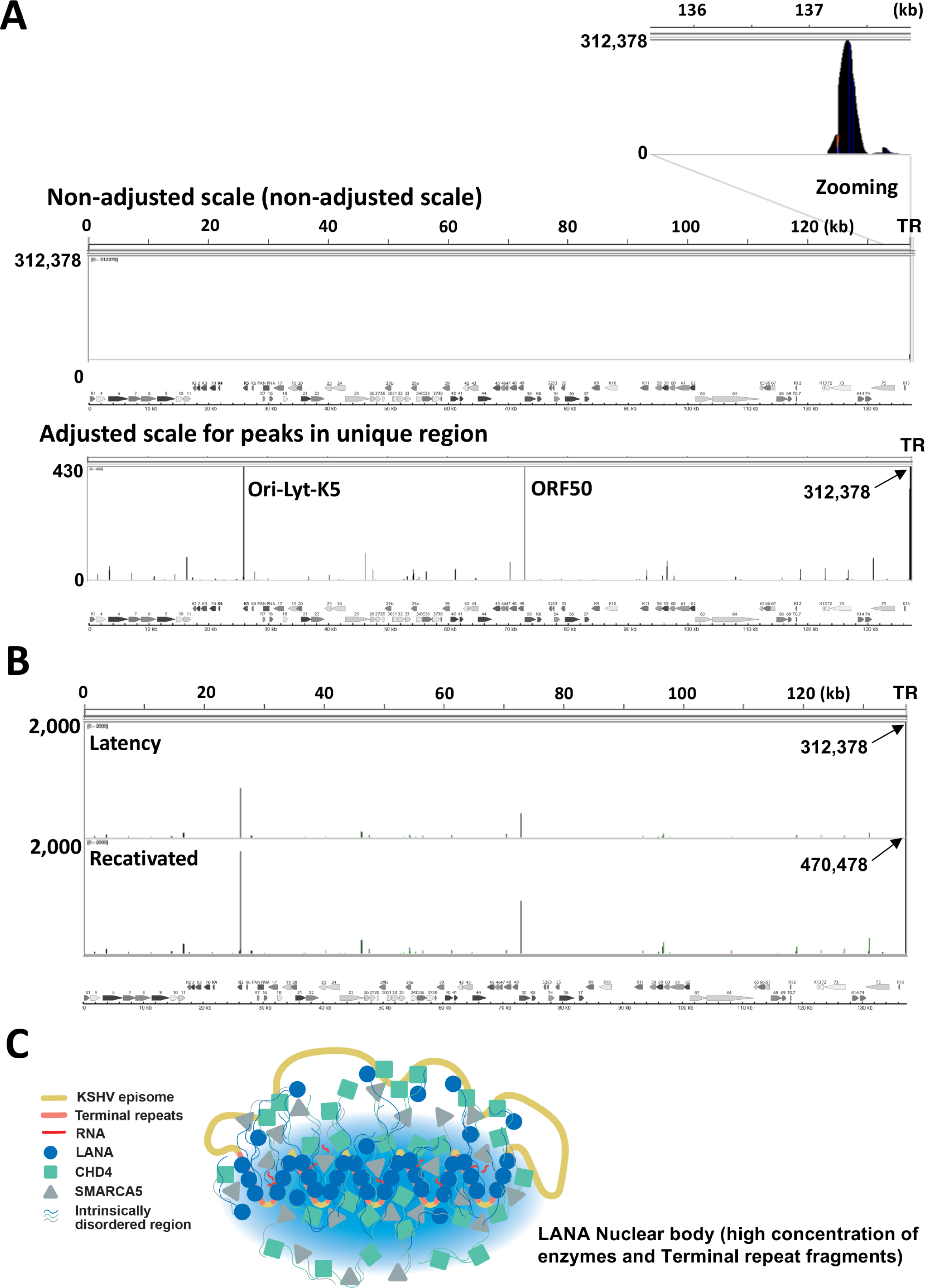
Terminal repeat does not form hard wire with unique region. The genomic looping with TR fragments was examined by isolating sequence reads from previous Hi-C data sets. The ligated DNA fragments with TR elements were aligned to the KSHV genome. The KSHV genome and ORF maps are shown at the bottom of the panel. **(A) Position of TR ligated DNA fragments.** Non-adjusted scale, which is based on the highest peaks (top panel), and the adjusted scale to show peaks in unique region (bottom panel) are presented. **(B) Changes in looping frequencies by reactivation.** Overall pattern of genomic loops was conserved but frequencies of genomic looping were increased by approximately 50%. **(C) TR serves as a genomic element to form flexible large protein storage with LANA nuclear bodies.** LANA is accumulated at TR region with direct DNA bindings and increases the concentration of LANA interacting proteins such as CHD4, SMARCA5, and BRD4. Locally concentrated enzymes support the regulation of lytic gene promoters with LANA (Fig. 3).

### TR supports LANA and K-Rta transcription function

The H3K27Ac histone modification and enrichment of transcription enzymes suggest that TR may possess transcription regulatory function, when it localizes proximity to promoter. To study this, we first cloned TR from KSHV BAC16 by homologous recombination with recombineering technique (39) (Fig. 5A). Homology arms are included in long primer sequence (**Table 1**), and the primers were used to amplify pBlueScript plasmid vector. The resulting PCR fragment containing plasmid DNA’s origin, ampicillin-resistant cassette, and partial multiple cloning sites, were transduced into BAC16 containing *E.coli*. The successful recombination generates circular plasmid in BAC containing *E.coli* which outgrowths via high copy origins of plasmid DNA replication. The ampicillin-resistant colonies were then screened for the presence of TR, and number of TR copies was examined by size of plasmid DNA after restriction enzyme digestion (Fig. 5B). To study association of increased TR copy numbers with transcription function, we isolated pBS-TR vector encoding 0, 2, 4, 6 copies of TRs [pBlueScript (BS)-TR]. We then cloned the luciferase reporter cassette into the pBS-TR vector. KSHV reporter libraries(26) generated with pGL3 basic vector was used as a PCR template. The reporter cassette includes synthetic poly(A) sites at upstream of the inserted promoter and SV40 poly(A) sites downstream of the luciferase gene (Fig. 5C). We cloned reporter constructs in two orientations (left and right: Fig. 5C) and examined effects on directionality. The results demonstrated that having TR fragments, KSHV PAN RNA promoter activity was strongly enhanced in independent of the orientation in 293FT cells (Fig. 5D). However, the TR-left PAN promoter, in which TR fragments are physically closer to PAN RNA promoter, showed higher luciferase activity. Having TR fragments strongly synergized K-Rta-mediated reporter activation with all promoter tested (Fig. 5E). On the other hand, while presence of TR increased basal promoter activity (TR0 vs TR4 with vector control in Fig. 5G), LANA seemed to acquire stronger gene repression function in the presence of TR. The effects were exaggerated when we deleted the LANA IDR domain, which may allow LANA to interact with other IDR-containing transcription factors, presumably other coactivators (Fig. 5G). The results suggested that TR fragments increased basal levels of transcription activity (TR0 vs. TR4) in majority of lytic gene promoters, and enhanced K-Rta transactivation function synergistically. Increasing the TR copy number further stimulated promoter activity, but the effects were not always linear in 293FT cells. The enhancement of promoter activity was saturated with 4 copies of TR assay, and promoter activity started to decline in presence in TR6 in 293FT cells (**S-**Fig. 2).

**Table 1.**
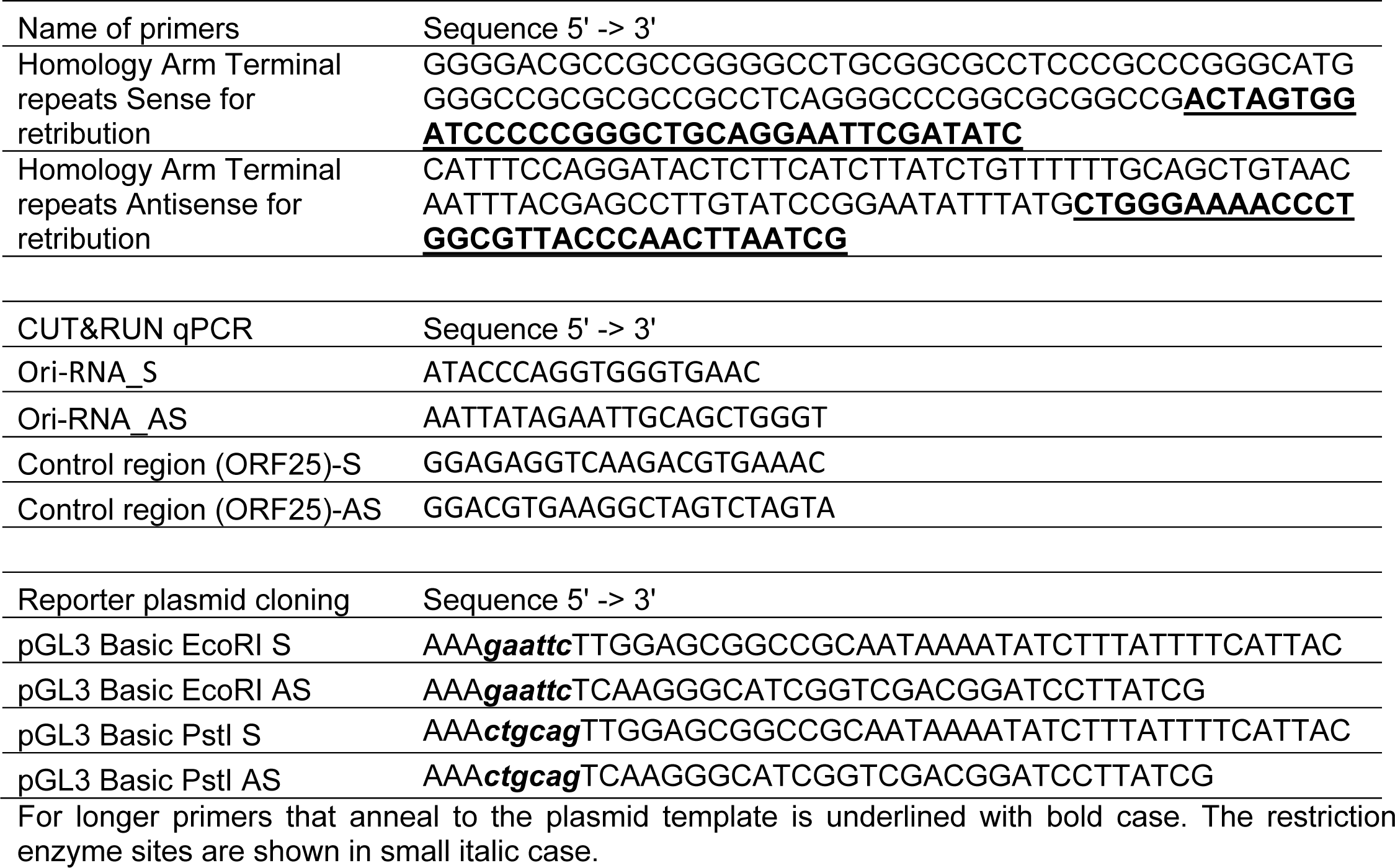
Primer and DNA fragments used in this study.

**Figure 5.**
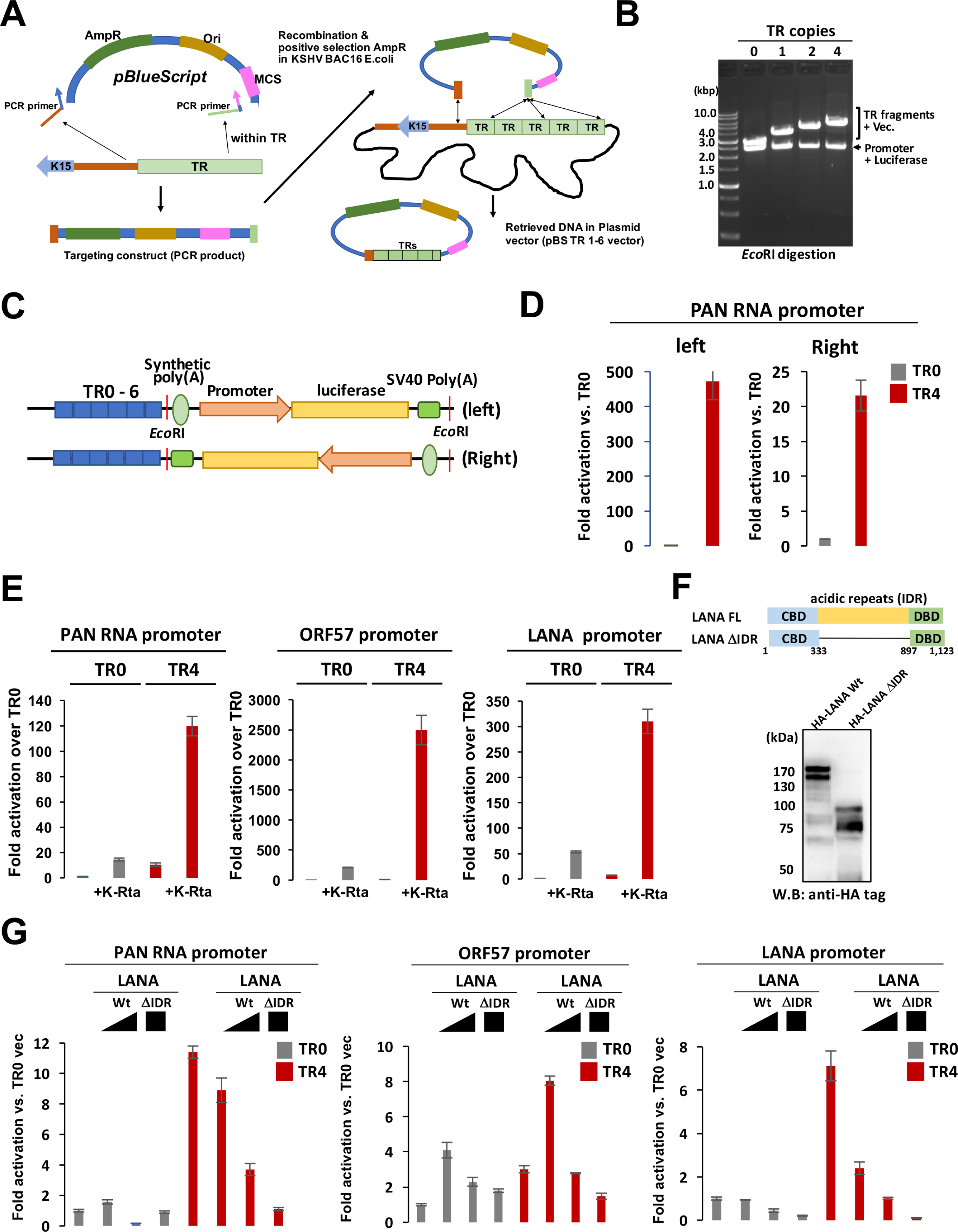
Terminal repeat fragment possesses enhancer function. (A) Recombination mediated TR cloning. Schematic diagram of TR fragment cloning. Long primers with homology arms to the end of the unique sequence and TR sequence was used to amplify the pBlueScript plasmid. The amplified DNA fragment was transformed into BAC16 containing GS1783 for recombination. **(B) Confirmation of TR insertion.** The EtBr stained agarose gel is shown. The copy number of TR was determined by an 801bp incremental size increase. **(C) Schematic diagram of the luciferase reporter construct.** Luciferase reporters were amplified from the pGL3 vector and cloned into *EcoR*I sites in two different orientations (left and right). **(D) TR activates PAN RNA promoter in orientation independent manner.** Reporter constructs were transfected into 293FT cells, and luciferase activity was measured 48 hours post transfection. Luciferase value with TR0 was normalized as 1, and fold activation is shown. **(E) TR enhances the K-Rta transactivation function.** Indicated reporter constructs (right direction) were co-transfected with K-Rta expression plasmid in 293FT cells. Fold activation over TR0 with vector control transfection is shown. **(F) Diagram and expression of LANA mutant.** The LANA acidic domain with acidic repeat domain deletion was constructed for reporter assay. The position of amino acid deletion and expression in 293FT was shown. **(G) TR enhances LANA’s transcription function.** Indicated reporter constructs (1 ug) were co-transfected with either full-length LANA expression plasmid (0.5 or 2.0 μg) or LANAΔIDR (2.0 μg). Luciferase values were measured 48 hours post transfection. For luciferase reporter assays, error bars with standard deviation for triplicated samples were shown. We performed reporter assays at least three times for each setting and representing results of one of the replicates are shown.

Finally, the enhancer activity was tested with KSHV infected 293FT cells. We first prepared infectious KSHV from BAC16-Wt iSLK cells, and generated KSHV-infected 293FT cells with hygromycin selection (Fig. 6A). The KSHV infection were confirmed with IFA and immunoblotting (Fig. 6A, B). The KSHV-infected 293FT cells were co-transfected with K-Rta expression plasmid with reporter constructs, and examined enhancer activity. The results again showed that TR enhanced K-Rta-mediated transcription function synergistically, perhaps even more so than in non KSHV infected 293FT cells (Fig. 6C). The results suggested that other viral proteins may facilitate promoter activation.

**Figure 6.**
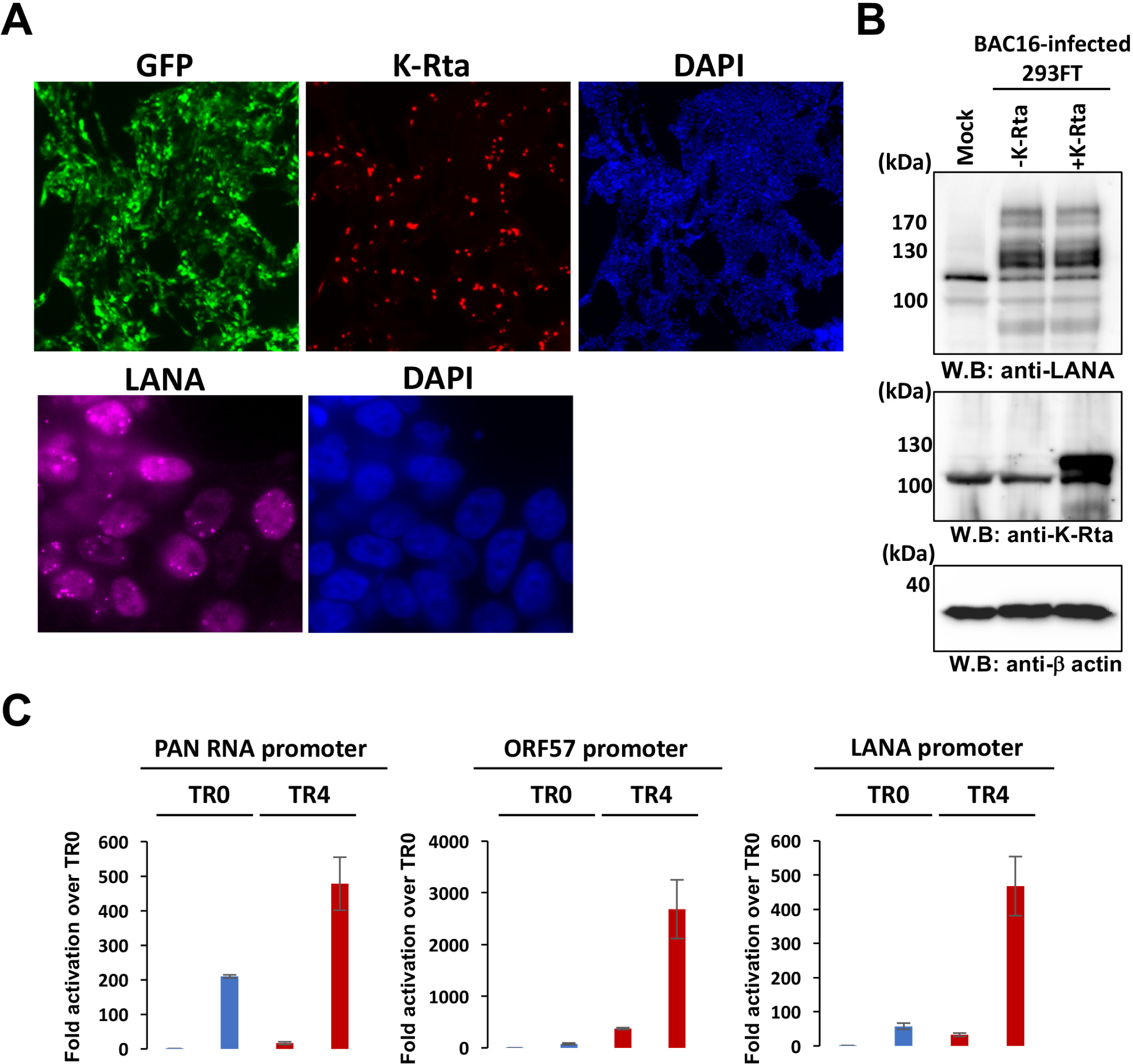
Terminal repeat fragment possesses enhancer function in KSHV-infected 293FT cells. **(A) Generation of KSHV-infected cells.** The BAC16 Wt KSHV was infected in 293FT cells and selected with hygromycin. The BAC16 stable 293FT cells were stained with the indicated antibody after transfection of the K-Rta expression plasmid. DNA was stained with DAPI. **(B) immunoblotting.** Infection and expression of K-Rta were confirmed by immunoblotting. Indicated antibodies were used for probing proteins. The β-actin served as loading control. **(C) Luciferase assay.** Indicated reporter constructs were transfected with either vector control or K-Rta expression plasmid. Luciferase values was measured after 48 hours transfection. Luciferase unit with TR0 reporter with vector control designated as 1, and fold activation is shown.

Taken together, we propose that KSHV TR acts as an "enhancer" for both transcription activation and repression with K-Rta and LANA, respectively. Our study suggests that KSHV evolved a very clever design of the KSHV "enhancer", which encodes arrays of high affinity LANA DNA binding domain to create phase separate LANA NB at TR, which positions LANA to control KSHV lytic gene promoters.

## Discussion

While KSHV promoter activity has been measured in isolated reporter plasmids and a number of key cellular transcription factors were identified as an important regulator for KSHV latency-lytic switch (26, 40–43), a mechanism through which many viral promoters are synchronously regulated remain unclear. This study aimed to provide an insight into the spatial and temporal KSHV gene regulation by focus on the transcription function of TR. The TR region takes up to one-fifth of limited KSHV genomic space(36, 44).

To identify critical regulatory domains of latent chromatin, we first established a histone modification map with CUT&RUN analyses(45). The histone modification predicts which genomic regions are accessible to DNA-binding proteins. The CTCF, SMC1A, Hi-C data also helped us to identify key genomic regions that formed a higher density of genomic hubs(23). Among active histone marks, H3K4me1, H3K4me3, and H3K27Ac are known to present at enhancers(46, 47). Subsequent studies showed that H3K4me1 is a marker for poised enhancers. When the poised enhancer is activated, the enhancer begins to possess H3K27Ac marks with or without H3K4me1(48). According to the definition, KSHV TR is an active enhancer, whose access is however restricted by LANA, presumably through a recruitment of CHD4 (24, 49).

We initially expected highly conserved H3K27me3 marks on KSHV latent chromatins, since latent chromatin is largely silenced. Studies by others elegantly demonstrated recruitment of PRC1/2 complex on KSHV genomes in iSLK cells (50–52). To our surprise, we found that degree and position of H3K27me3 marks were varied among naturally-infected three PEL cell lines (Fig. 1B) and also experimentally infected iSLK cells. Compared with BC3 cells, amount of H3K27me3 in BC1 at late gene cluster were much less and the position of the histone marks also varied among cell lines. Considering each PEL cell harbors 30+ episomes and studies have been performed with cell population, we think that a selected fraction of infected episomes may possess H3K27me3 modifications, yet KSHV episomes are not actively transcribed in PEL cells. The results may indicate that pausing RNAII elongation at the unique region but not global chromatin condensation(s) may be the major contributor for the lytic gene silencing. On the other hand, active histone modifications such as H3K27Ac and H3K4me3 were more conserved at specific genomic regions in PEL cell lines and experimentally-infected cells. This suggests that maintaining specific active genomic domain is the strategy, which KSHV has evolved for establishing inducible chromatin in presence of K-Rta. Importantly, these active regions possess the major K-Rta and LANA binding sites, and the K-Rta binding sites precisely positioned next to the LANA complex tethered nucleosome. For that, we speculate that very high LANA protein concentration at TR may help to ensure LANA and LANA interacting protein recruitment at the LANA binding sites in unique region. Although more mechanistic studies have to be followed, recruitment of SWI/SNF via K-Rta DNA binding is expected to evict the nucleosome, hence LANA complex from the Ori Lyt DNA. Indeed, earlier study showed that establishing nucleosome free loci at K-Rta promoter regions during reactivation (53). Together, we propose that eviction of LANA-tethered nucleosome at Ori-Lyt, where RNAPII paused, is a well-designed and highly efficient KSHV Latency-Lytic switch mechanism.

A combination of previous proteomics studies on LANA and K-Rta allowed us to draw the putative protein interaction maps (Fig. 3). We showed that cellular proteins that interact with LANA in latently infected cells may also accommodate enzymes that interact with RNAPII and K-Rta during reactivation. Multiple copies of TR recruit a significant amount of LANA and BRD4. Those two proteins can flexibly harbor many other histone enzymes as protein hubs. We speculate that large and highly disordered acidic repeat regions (IDR) of LANA should play an important role in capturing such enzymes at TR. Having a larger IDR would increase efficacies for trapping another IDR containing proteins dynamically. The mechanism is analogous to a system operated at cellular super enhancer with interaction between two IDR containing proteins, BRD4 and MED1(11). Consistent with this, LANA became a stronger repressor when we deleted the IDR domain (Fig. 5G). Importantly, KSHV K-Rta interacts with MED1 during reactivation (Fig. 3), and KSHV LANA associates with BRD4 at TR and Ori-Lyt region. Taken together, TR is a well-poised enhancer for KSHV lytic gene promoters, and viral gene promoters are activated by recruitment of MED1, a component of the mediator complex, via K-Rta, that should create many inducible genomic loops among BRD4 bound regions (TRs and RNAPII poised regions in unique region). Importantly, we demonstrated such genomic looping shift during reactivation, in which KSHV 3D genomic structure was more compacted by the generation of larger transcription regulatory domains, and 3D genomic structures became squeezed doughnuts like shape (23).

Suppose the activity of KSHV TR is essential for lytic gene expression. How is the KSHV TR initially activated during *de novo* infection and maintained to be activated in the host cells? Does constitutively active NF-κB and STAT pathways frequently seen in KSHV infected cancer cell lines contribute to the active episome maintenance? From an evolutional point of view, we speculate that KSHV vGPCR, vIRFs, or vIL-6 may have additional functions to ensure establishment of re-activatable latent chromatin. We also think large LANA NBs are like a "solar panel" to extract energy by intercepting cellular signaling events via highly concentrated LANA/BRD4 complex at TR. The highly-concentrated IDR proteins would absorb transcription enzymes from nearing cellular transcription factors because affinity of interaction among cellular transcription factors and enzymes (i.e., SWI/SNF) are relatively weak in order to share them among other cellular transcription factors for dose-dependent regulation, except viral transcription factor like K-Rta (54). TR fragments and unstructured acidic repeats in LANA protein seems to be designed to concentrate the protein complex near the KSHV unique protein coding regions. Consistent with this, we have seen number of transcription factors (e.g., STAT3, p65, IRF4, BATF), coactivator enzymes (SMARCA4, SMARCC1), DNA damage related enzymes, SUMO enzymes, and many other proteins that form dynamic and flexible protein complexes, co-localize with the LANA NBs in BCBL-1. While these ideas await further examination, TRs may, in return, function like a sponge to weaken the ability for cellular TFs to directly regulate KSHV reactivation, which should help to maintain latent state for immune evasion. This is partly see in reporter assays, in which the TR6 but not TR0 reporter could repress promoter activation, when we overexpressed LANA. We also found that recombinant KSHV with only 2 copies of TR showed highly lytic phenotypes with frequent spontaneous reactivation compared with 10 copies of TR (unpublished observation).

If the LANA is highly concentrated at proximity to lytic gene promoters, how KSHV dissolve such condensates for continuing transcription burst by preventing from rebinding? An exciting study was recently published by Dr. Roger A. Young’s laboratory (55). The study demonstrated that the amount of transcribing RNA is critically associated with its role in recruitment and dissolving protein condensates at enhancer and promoter regions, thereby regulating enhancer-promoter interactions by modulating the concentrations of coactivator complexes at the genomic sites. The authors demonstrated that high levels of RNA from robust gene transcription bursts can dissolve condensates; this mechanism generates a negative feedback loop (55). The authors also showed that charge balance of electrostatic interactions is, in part, responsible for the transcription-mediated feedback regulation (55). Consistent with the study, we previously showed that CHD4 and LANA are sequestered away from the PAN RNA promoter when we triggered KSHV reactivation (24, 56). We also demonstrated that CHD4 is a sequence-independent RNA binding protein, and CHD4 is a part of protein condensates with LANA (24). Therefore, robust PAN RNA expression (>10^5^ copies/cell during reactivation(57)) may be another clever design to dissolve LANA condensates at LANA recruitment sites (e.g., Ori-Lyts and K-Rta promoter). Two unique genomic properties, TR copy number and exceptional viral lncRNAs expression, would establish tighter control for the KSHV latency-lytic switch, which should increase the stability of maintaining inducible latent chromatins.

In summary, we suggest a KSHV latency-lytic switch model based on KSHV 3D genomic structure, K-Rta and LANA interacting protein recruitment sites, and chromatin modification map. Regulation of dynamic protein interaction at Ori-lyt during reactivation, and molecular mechanism(s) of maintenance of epigenetically active status of the viral super enhancer should provide an important target to intercept KSHV replication cycles.

## MATERIALS AND METHODS

### Chemicals, reagents and antibodies

Dulbecco’s modified minimal essential medium (DMEM), fetal bovine serum (FBS), phosphate-buffered saline (PBS), Trypsin-EDTA solution, 100 X penicillin–streptomycin–L-glutamine solution were purchased from Thermo Fisher (Waltham, MA). Puromycin and G418 solution were obtained from InvivoGen (San Diego, CA). Hygromycin B solution was purchased from Enzo Life Science (Farmingdale, NY). The following antibodies were used for CUT & RUN, immunoblotting and flow cytometry: rabbit anti-BRD4 (Cell Signaling, E2A7X), rabbit anti-RNAPII (Millipore, clone CTD4H8), rabbit anti-H3K27ac (CST, clone D5E4), rabbit anti-H3K4me3 (Cell Signaling, clone C42D8), mouse anti-β-actin (Santa Cruz 47778), rabbit anti-gp130 (CST), rabbit IgG (CST, clone DA1E).

### Cell culture

293T cells were grown in DMEM containing 10% FBS and 1xPen-Strep-L-Gln at 37°C with air containing 5% carbon dioxide. BAC16 Wt stable iSLK cells were maintained in DMEM containing 10% FBS, 400 μg/ml hygromycin B, and 250 μg/ml G418, and 1xPen-Strep-L-Gln at 37°C with air containing 5% carbon dioxide. The KSHV BAC16-Wt infected 293FT cells were established after infection for BAC16-Wt virus with hygromycin selection (1,000 μg/mL) for 2 weeks.

### Preparation of pTR-reporter construct

KSHV TR fragments were cloned by adapting the recombineering technique (39). Homology arms that one targets unique region and another one targeting within the TR region were designed and used to amplify pBlueScript plasmid fragment. The linear PCR fragment that encodes portion of multiple cloning sites, the origin of plasmid replication, and ampicillin-resistant cassette, were purified from DNA agarose gel. Purified DNA fragments were then introduced to BAC16 containing GS1783 *E.coli* for red recombination(58). The ampicillin-resistant colonies were isolated and cultured in 32C for overnight. Outgrowing plasmids were purified with Qigen Mini-prep kits. Purified plasmids were used for restriction enzyme digestions to identify number of TR copies cloned. The pBluescript plasmid containing TR2, 4, or 6 was then used to transform into stbl3 strain (Invitrogen) for further plasmid amplification.

Luciferease reporter fragments were amplified from pGL3 KSHV promoter library with primers listed in Table 1. Amplified reporter DNA fragment was cloned into pBS-TR multiple cloning sites (*Eco*RI, *Pst*I, or *Hinc*II). The luciferase reporter fragments were cloned for two different orientations. Schematic diagram was also presented in a figure. The expression plasmid for pcDNA HA-K-Rta and pcDNA HA-LANA were described previously (59, 60). The position of cloned KSHV promoter fragments is listed previously (26). For generation of HA-LANA IDR deletion expression plasmid (pcDNA HA-LANAΔIDR), the DNA fragment was synthesized (IDTDNA), and cloned into pcDNA HA-vector. Synthesized DNA fragment was listed in Table 1.

### Cleavage under targets and release using nuclease (CUT&RUN)

CUT&RUN (45) was performed essentially by following the online protocol developed by Dr. Henikoff’s lab with a few modifications. Cells were washed with PBS and wash buffer [20 mM HEPES-KOH pH 7.5, 150 mM NaCl, 0.5 mM Spermidine (Sigma, S2626), and proteinase inhibitor (Roche)]. After removing the wash buffer, cells were captured on magnetic concanavalin A (ConA) beads (Polysciences, PA, USA) in the presence of CaCl_2_. Beads/cells complexes were washed three times with digitonin wash buffer (0.02% digitonin, 20 mM HEPES-KOH pH 7.5, 150 mM NaCl, 0.5 mM Spermidine and 1x proteinase inhibitor), aliquoted, and incubated with specific antibodies (1:50) in 250 μL volume at 4℃ overnight. After incubation, the unbound antibody was removed with digitonin wash buffer three times. Beads were then incubated with recombinant Protein A/G–Micrococcal Nuclease (pAG-MNase), which was purified from *E.coli* in 250 μl digitonin wash buffer at 0.5 μg/mL final concentration for 1 hour at 4 °C with rotation. Unbound pAG-MNase was removed by washing with digitonin wash buffer three times. Pre-chilled digitonin wash buffer containing 2 mM CaCl_2_ (200 μL) was added to the beads and incubated on ice for 30 min. The pAG-MNase digestion was halted by the addition of 200 μl 2× STOP solution (340 mM NaCl, 20 mM EDTA, 4 mM EGTA, 50 μg/ml RNase A, 50 μg/ml glycogen). The beads were incubated with shaking at 37 °C for 10 min in a tube shaker at 300 rpm to release digested DNA fragments from the insoluble nuclear chromatin. The supernatant was then collected by removing the magnetic beads. DNA in the supernatant was purified using the NucleoSpin Gel & PCR kit (Takara Bio, Kusatsu, Shiga, Japan). Sequencing libraries were prepared from 3 ng DNA with the Kapa HyperPrep Kit (Roche) according to the manufacturer’s standard protocol. Libraries were multiplex sequenced (2 × 150 bp, paired-end) on an Illumina NovaSeq 6000 system to yield ∼15 million mapped reads per sample.

CUT&RUN sequence reads were processed with fastp(61) and aligned to the human GRCh38/hg38 and KSHV reference genome (NC_009333.1) with Bowtie2 and yielding mapped reads in BAM files(62).

For CUT&RUN qPCR studies, TREx-K-Rta BCBL-1 cells were induced for lytic gene expression by incubating with doxycycline (1 ug/mL) and TPA (20 ng/mL) for 48 hours. TREx-BCBL-1 cells induced or non-induced was used for CUT&RUN and amount of released DNA fragments were examined by qPCR with primers listed in Table 1.

FASTQ files for the Hi-C experiments were processed through the HiCUP (v0.7.4) pipeline(63) using the KSHV reference genome (NC_009333.1). In the pipeline, Bowtie 2 (version 2.3.5.1) was used to map valid Hi-C ditags with a single restriction fragment, allowing only unique high-quality alignments across the genome. Invalid Hi-C ditags, such as dangling ends and PCR duplicates, were removed by the HiCUP Filter script. Valid Hi-C ditags were aligned to the TR region of the KSHV genome with Bowtie2(62) with al-conc option setting. Each forward and reverse read was independently mapped to the KSHV genome. The extracted paired reads were mapped again to the KSHV reference genome to identify three-dimensional genomic interaction with TR. The mapped reads were visualized with Integrative Genomics Viewer(64).

### RT-qPCR

Total RNA was extracted using the Quick-RNA miniprep kit (Zymo Research, Irvine, CA, USA). A total of 1 μg of RNA was incubated with DNase I for 15min and reverse transcribed with the High Capacity cDNA Reverse Transcription Kit (Thermo Fisher, Waltham, MA USA). The resulting cDNA was used for qPCR. SYBR Green Universal master mix (Bio-Rad) was used for qPCR according to the manufacturer’s instructions. Each sample was normalized to 18S ribosomal RNA, and the ddCt fold change method was used to calculate relative quantification. All reactions were run in triplicate. Primer sequences used for qRT-PCR are provided in the **Table. S1.**

#### STRING protein interaction visualization

Names of previously identified LANA neighboring protein with proximity biotin labeling (24) and K-Rta interacting complex (54), which is recruited to RNAPII during reactivation were collected and visualized with STRING, a public database of known and predicted protein-protein interactions. Dynamic protein interaction among K-Rta and LANA protein complex during reactivation was also visualized with STRING after manually combine proteins names together. The list of proteins based on previous analyses (p<0.05) with chromatin regulatory function are all included in this analysis.

### Luciferase assay

HEK293FT or KSHVr2.19-infected HEK293FTcells were seeded onto 12-well plates at 2.0 x 10^5^/well. The cells were transfected with expression plasmid encoding the HA-epitope tagged K-Rta, LANA, or together, along with pBS-TR reporter constructs. The TR reporter constructs encodes varied number of TR repeats and KSHV gene promoter, which is cloned in front of luciferease coding sequence. Cell lysates were prepared with 1% TritonX-100 and 0.5% NP-40 in PBS 48 hours after transfection. Luciferase activity was measured according to the manufacturer’s protocol by using Varioskan LUX (Thermo Scientific). At least three independent measurements were performed for each setting.

### Immunoblotting

Protein lysates from luciferase assays were subjected to 8% SDS-PAGE gel and transferred to PVDF membranes (Millipore-Sigma, St. Louis, MO, USA). Membranes were incubated with 5% nonfat milk at room temperature for 15 minutes for blocking, and then incubated with the primary antibody at 4°C overnight. After washing membrane with TBST three times, the membrane was then incubated with horseradish-peroxidase-conjugated secondary antibody (Santa Cruz) at RT for 1 hour. The final dilution of the primary antibody was 1:2,000 for anti-HA tag (Covance) and 1:3000 for anti-β-actin (Millipore-Sigma).and K-Rta (65).

### Statistical analysis

Experimental replicates of at least 3 for each sample, including negative controls, were prepared whenever applicable. Results are shown as mean ± SD from at least three independent experiments. Statistical analyses were performed using GraphPad Prism 9.4.1 software. A value of p < 0.05 was considered statistically significant.

## Data availability

The CUT and RUN data were deposited in the NCBI Gene Expression Omnibus (GEO) database under accession number GSE241949.

## Acknowledgments

We want to thank Drs. Ken-ichi Nakajima, Somayeh Komaki, Kazushi Nakano, and Kang-Hsin Wang for technical support. We also like to thank UC Davis Comprehensive Cancer Center Genomic Shared Resource members for depositing sequence data.

This research was supported by public health grants from the National Cancer Institute (CA225266, CA232845) and the National Institute of Allergy and Infectious Disease (AI167663) to Y.I. The Genomics is supported by the UC Davis Comprehensive Cancer Center Support Grant (CCSG) awarded by the National Cancer Institute (NCI P30CA093373).

